# Targeted Deletions Uncover Previously Uncharacterised C_1_-fixing Cluster Genes as Essential for Autotrophy in *Clostridium autoethanogenum*

**DOI:** 10.64898/2026.11.07.765054

**Authors:** Ugochi J. Nwaokorie, Mariia Vorontsova, Kurshedaktar M. Shaikh, Kristina Reinmets, Kaspar Valgepea

## Abstract

Recycling carbon oxides from industrial off-gases and gasified organic waste using gas fermentation offers a sustainable route for biomanufacturing of fuels and chemicals. Acetogens are attractive biocatalysts here and fix CO and CO_2_ via the Wood-Ljungdahl pathway (WLP). However, the incomplete functional annotation of the associated C_1_-fixing gene cluster limits rational engineering of acetogen cell factories. Here, we investigated five highly conserved but previously uncharacterised C_1_-fixing cluster genes to quantify their contributions to the growth of the model-acetogen *Clostridium autoethanogenum* using targeted CRISPR/Cas9-mediated single-gene deletions, physiological characterisation, comparative computational structural analyses, CRISPRi, and functional genomics. Importantly, we identified *acsV*, *lpdA*, *cooC,* and *cooC2* as essential for autotrophy but not heterotrophy, whereas deletion of *gcvH* reduced autotrophic growth and shifted carbon flux from acetate towards 2,3-butanediol. Amino acid supplementation experiments and computational analyses indicate that *lpdA* and *gcvH* do not primarily function in glycine or serine biosynthesis but instead fulfil specialised roles in WLP-associated C_1_ and redox metabolism. Furthermore, genetic and physiological analyses showed that the two CooC-family ATPases, CooC and CooC2, have non-redundant roles in growth, with only *cooC2* deletion impairing heterotrophy. Acetate supplementation tests revealed that Δ*acsV* cells retained metabolic activity despite abolished autotrophy, and computational analyses suggest a likely role for *acsV* in ATP-dependent corrinoid activation. Surprisingly, CRISPRi-mediated repression of *acsV* was translationally buffered, resulting in altered carbon partitioning without impairing autotrophy. Together, our findings provide insights into the physiological importance of the C_1_-fixing cluster genes in acetogens and support their metabolic engineering as gas-fermenting cell factories.

## Introduction

Industrial waste gases containing one-carbon (C_1_) compounds (CO, CO_2_, CH_4_) contribute significantly to global anthropogenic carbon emissions, but represent an abundant, yet underutilised resource. Gas fermentation uses specialised chemolithoautotrophic microbes – such as acetogens, hydrogenotrophs, and methanotrophs – to convert these C_1_ gas streams into valuable biofuels, chemicals, and feed protein, providing a promising route for sustainable carbon utilisation and circular value creation.^1,2^ Distinct carbon fixation and energy conservation pathways in these microbes enable the conversion of CO-, CO_2_-, and CH_4_-rich gases into biomass and value-added products.^3,4^ Among these, acetogens are the most advanced gas-fermenting biocatalysts, with *Clostridium autoethanogenum* deployed at an industrial scale for ethanol production and at pilot-scale for acetone and isopropanol production from carbon oxides. ^5–7^ Although the biochemistry of the ancient Wood-Ljungdahl pathway (WLP) that acetogens use for fixing carbon oxides is well characterised,^8–10^ the incomplete understanding of acetogen metabolism continues to limit rational strain engineering to advance their use as industrial cell factories for gas fermentation.^11^ Until recently, the labour-intensive development of efficient gene-editing tools constrained both functional characterisation and metabolic engineering of acetogens.^11,12^ Thus, the functions of several conserved genes within the C_1_-fixing gene cluster harbouring the WLP genes remain unresolved, and it is unknown whether these genes affect autotrophic growth, heterotrophic growth, or both.^13^

The five genes with unclear functions in the C_1_-fixing gene cluster in the *C. autoethanogenum* LAbrini strain,^14^ are computationally annotated as: 1) corrinoid activation/regeneration protein (*acsV*; LABRINI_07955, following gene IDs stated without ‘LABRINI_’); 2) glycine cleavage system H protein (*gcvH;* 07960); 3) AAA family ATPase (*cooC;* 07985); 4) dihydrolipoyl dehydrogenase (*lpdA;* 07990); and 5) carbon monoxide dehydrogenase accessory protein (*cooC2;* 08020) (Figure 1).

**Figure 1.**
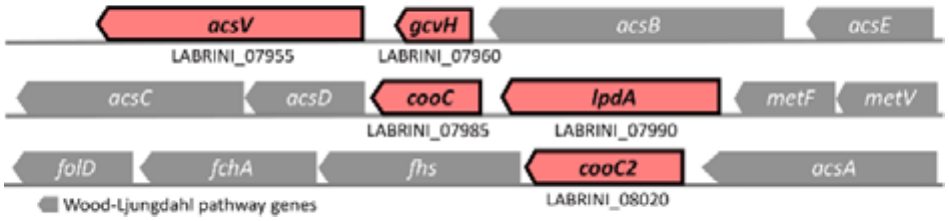
C_1_-fixing gene cluster of *C. autoethanogenum* LAbrini. The gene cluster contains 15 genes, including five genes with unconfirmed biochemical functions (coloured light red; LABRINI_07955; 07960; 07985; 07990; 08020):^13^ *acsV* - corrinoid activation/regeneration protein; *gcvH* - glycine cleavage system H protein; *cooC* - AAA family ATPase; *lpdA* - dihydrolipoyl dehydrogenase; *cooC2* - carbon monoxide dehydrogenase accessory protein. See Table S4 of a previous study for descriptions of Wood-Ljungdahl pathway genes.^15^

All five genes are highly conserved in the C_1_-fixing gene clusters across acetogens,^16,17^ and exhibit high transcript and protein expression levels similar to cluster members of WLP genes.^15,18–22^ This may indicate that the entire cluster is likely expressed as a single polycistronic mRNA, as no transcription initiation or termination sites are detected within the cluster in LAbrini.^23^ The co-expression of these five genes, despite the energetic cost of their high protein expression, suggests that they may contribute to autotrophic growth of acetogens.

Three of the five gene products (AcsV, CooC, and CooC2) are annotated as proteins linked to the bifunctional nickel (Ni)-dependent carbon monoxide dehydrogenase/acetyl-CoA synthase (CODH/ACS) enzyme complex that is essential for autotrophic growth of *C. autoethanogenum* and whose assembly and protein interactions have been extensively characterised.^24–28^ Both *cooC* and *cooC2* are proposed to belong to the Min-D type ATPases involved in ATP-dependent insertion of Ni into CODH/ACS.^29^ Interestingly, studies of the related C_1_-fixing microbe *Carboxydothermus hydrogenoformans*, which also uses the WLP, show that CooC-family MinD-type maturases can share sequence and structural similarities but interact differently with the CODH/ACS complex, suggesting distinct functional roles.^30,31^ For example, the CooC-type and AcsF-type subgroups of the CooC family have been linked to Ni insertion into the CODH active site and formation of the ACS bimetallic A-cluster, respectively.^30^ The *acsV* gene was previously annotated as a ferredoxin in *C. autoethanogenum*^32^ but has recently been re-annotated to putatively encode a corrinoid activation/regeneration protein associated with the CODH/ACS complex.^33^ Furthermore, there has been considerable debate regarding the presence and functions of the remaining two genes, *lpdA* and *gcvH*, within the C_1_-fixing gene cluster.^34^ Both genes are annotated as components of the glycine cleavage system (GCS), a central part of the reductive glycine cleavage pathway (rGlyP), which has been proposed to support the WLP in CO_2_ fixation by linking glycine and serine metabolism with tetrahydrofolate (THF)-dependent C_1_ metabolism.^16^ Additional hypotheses suggest that *lpdA* and *gcvH* may contribute more directly to WLP-associated redox metabolism, including acting as a potential electron donor for the methylene-THF reductase.^35^

Here, we used CRISPR/Cas9-based single-gene deletions to investigate the functions of the five aforementioned conserved but previously uncharacterised genes within the C_1_-fixing gene cluster of the model-acetogen *C. autoethanogenum*. We identify four of these genes as essential for autotrophic growth. We further demonstrate that the WLP-associated *gcvH* fulfils a specialised, non-essential role in autotrophic metabolism, and that the two proposed CooC-family MinD-type ATPases indeed perform non-redundant functions during autotrophic growth, while computational analysis supports a likely role for *acsV* in ATP-dependent corrinoid activation. Together, our findings resolve key physiological requirements of genes within the highly conserved C_1_-fixing gene cluster in acetogens and provide new insight into its functional organisation for autotrophy.

## Results and Discussion

### 3.1 Deletions of C_1_-fixing gene cluster genes with unclear functions affect heterotrophic growth and by-product profiles

To investigate the roles of the five genes with unclear functions in the C_1_-fixing gene cluster of *C. autoethanogenum* LAbrini – *acsV*, *gcvH*, *cooC*, *lpdA*, and *cooC2* (Figure 1) – CRISPR/nCas9 plasmids were constructed (Figure 2A) to generate single-gene deletion strains for each gene using a previously described workflow.^19^ Sanger sequencing confirmed the clean deletion of each target gene (Figure S1), validating our targeted gene-deletion workflow in a difficult-to-engineer acetogen.

**Figure 2.**
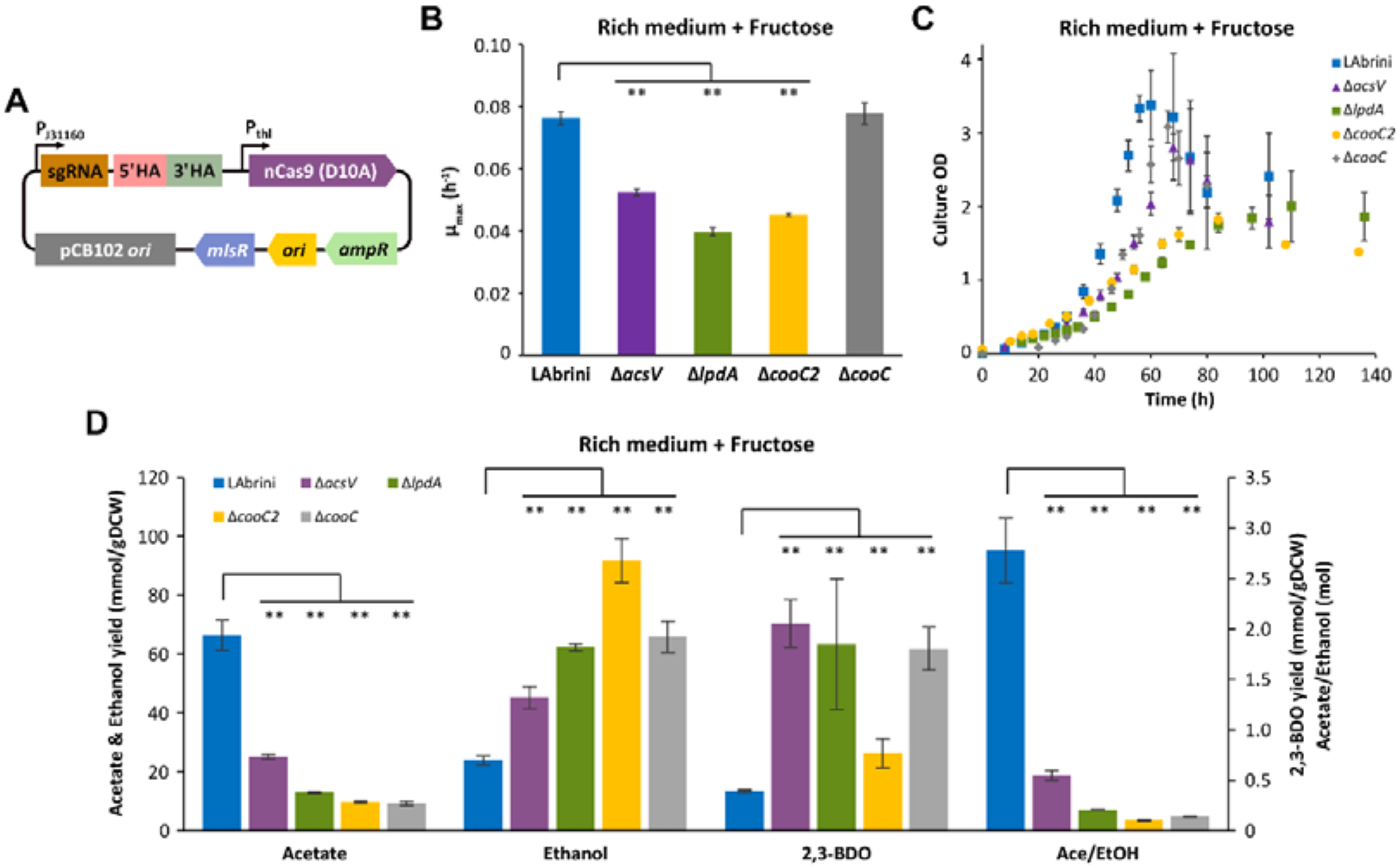
Heterotrophic characterisation of *C. autoethanogenum* LAbrini, Δ*acsV*, Δ*lpdA*, Δ*cooC2*, and Δ*cooC* on fructose in PETC-MES medium with YE (rich medium) in bottle batch cultures. (A) CRISPR/nCas9 construct used for targeted single-gene deletions. 5’HA and 3’HA indicate upstream and downstream homologous arms (HA) for gene deletion, respectively. (B) Maximum specific growth rates (µ_max_). (C) Growth profiles. (D) Growth by-product yields (mmol/gDCW) and acetate-to-ethanol ratios (Ace/EtOH; Cmol). Data represent mean ± standard deviation between three biological replicates. Asterisk denotes values statistically different according to two-tailed unpaired *t*-test (**, p < 0.01). LAbrini, *C. autoethanogenum* LAbrini strain; HA, homology arm; OD, optical density; sgRNA, single guide RNA; YE, yeast extract; gDCW, gram of dry cell weight; 2,3-BDO, 2,3-butanediol.

We first characterised the effects of gene deletions on heterotrophy by growing four strains – Δ*acsV,* Δ*lpdA,* Δ*cooC2*, and Δ*cooC* – in bottle batch cultures using PETC-MES medium with yeast extract (YE; hereafter, rich medium) and fructose. While no difference in maximum specific growth rate (µ_max_) was observed for Δ*cooC* (0.08 ± 0.003; average ± standard deviation between biological replicates) compared to LAbrini (0.08 ± 0.002), significantly lower values were observed for the other three characterised deletion strains (Figure 2B-C).

Specifically, Δ*acsV,* Δ*lpdA*, *and* Δ*cooC2* showed µ_max_ values of 0.05 ± 0.001, 0.04 ± 0.001, and 0.05 ± 0.001, which were 31, 48, and 41% lower than LAbrini, respectively (Figure 2B). Additionally, the maximum culture optical density (OD_max_) was significantly lower for Δ*acsV* (2.80), Δ*lpdA* (2.01), and Δ*cooC2* (1.44) compared to LAbrini (3.4) (Figure 2C), showing that these three genes clearly contribute to heterotrophic growth of *C. autoethanogenum*.

Analysis of growth by-product production during exponential growth revealed a significant reduction in the acetate yield (mmol/gram of dry cell weight [gDCW]) for all strains (Δ*acsV*: 25.1 ± 0.8; Δ*lpdA*: 13.1 ± 0.2; Δ*cooC2*: 9.8 ± 0.1; Δ*cooC*: 9.3 ± 0.7 mmol/gDCW) relative to LAbrini (66.5 ± 5 mmol/gDCW) (Figure 2D). In contrast, the ethanol yield was significantly higher for all strains (Δ*acsV*: 45.4 ± 3.7; Δ*lpdA*: 62.4 ± 1.2; Δ*cooC2*: 91.8 ± 7.5; Δ*cooC*: 65.9 ± 5.3 mmol/gDCW) compared to LAbrini (23.9 ± 1.5 mmol/gDCW). These changes thus translated into significantly lower acetate-to-ethanol ratios (Ace/EtOH; Cmol) for the four deletion strains (Figure 2D). Similarly to ethanol, increased yields for another reduced growth by-product – 2,3-butanediol (2,3-BDO) – were observed in the four strains: Δ*acsV*: 2.06 ± 0.24; Δ*lpdA*: 1.85 ± 0.65; Δ*cooC2:* 0.77 ± 0.14; Δ*cooC:* 1.81 ± 0.21 vs LAbrini: 0.39 ± 0.01 mmol/gDCW. Notably, while there was no growth impairment for Δ*cooC* (Figure 2B-C), product yields were similarly affected as with other deletion strains.

The pronounced decrease in acetate production, accompanied by increased ethanol and 2,3-BDO production across the four strains, shows substantial redistribution of carbon and reducing equivalents in *C. autoethanogenum* following deletion of any of the four genes. As the WLP plays a central role in maintaining redox balance in acetogens,^9^ these results suggest that *acsV*, *cooC*, *lpdA*, and *cooC2* genes contribute to optimal WLP function, with their deletion constraining electron flow and forcing cells to redirect excess reducing power toward the more reduced growth by-products, ethanol and 2,3-BDO, which act as redox sinks.^36,37^ While this metabolic shift may restore redox homeostasis, it could come at the expense of ATP due to the lower ATP gain from these reduced products vs acetate, potentially explaining the slower growth of Δ*acsV*, Δ*lpdA*, and Δ*cooC2*. Lower ATP production may also arise from reduced net acetate formation, for instance, if ethanol production shifts from the ATP-coupled, ferredoxin (Fd)-consuming aldehyde:Fd oxidoreductase (AOR) route to the bifunctional aldehyde/alcohol dehydrogenase (AdhE) route.

A previous gene essentiality analysis using the transposon directed insertion-site sequencing (TraDIS) approach in the *C. autoethanogenum* wild-type JA1-1 strain reported that both *cooC* and *lpdA* are essential for heterotrophic growth on YTF medium, while the other genes – *acsV*, *cooC2*, and *gcvH* are dispensable.^38^ We, however, believe that deletion of the genes investigated in this work is not expected to be lethal for heterotrophic growth of LAbrini, as deletion of the key WLP enzyme AcsA (CODH) in *C. autoethanogenum* JA1-1 results in similarly impaired growth and reduced acetate production.^39^ This supports the view that, while the WLP contributes to redox balancing and carbon flux during growth on sugars, it is not essential for heterotrophic growth, explaining the non-lethal but impaired phenotypes observed here.

### 3.2. Autotrophic characterisation reveals distinct physiological requirements of glycine cleavage–associated *gcvH* and *lpdA* genes

We next investigated the effects of deleting each of the five genes during autotrophy of *C. autoethanogenum*, guided by existing computational annotations of their proposed gene products (Figure 1). All autotrophic growth tests were performed using rich medium under autotrophic syngas (CO+CO_2_+H_2_) conditions, unless noted otherwise.

We first examined the Δ*lpdA* and Δ*gcvH* strains during autotrophy, where both deleted genes are proposed to be involved in the GCS, a central carbon pathway widely distributed across eukaryotes, archaea, and bacteria, including acetogens.^40^ Deletion of *gcvH* caused 22% slower growth (0.072 ± 0.001 vs 0.093 ± 0.002) (Figure 3A) and 32% lower OD_max_ (0.88 ± 0.04 vs 1.30 ± 0.04) compared to LAbrini (Figure 3B). Because *gcvH* deletion did not abolish growth under the more metabolically restrictive autotrophic conditions, the gene is unlikely to be essential for heterotrophic growth. Notably, Δ*gcvH* showed a 71% lower acetate yield (47.5 ± 11.6 vs 164.5 ± 9.9 mmol/gDCW) with a 7-fold elevated 2,3-BDO yield (6.4 ± 1.3 vs 0.87 ± 0.08 mmol/gDCW) compared to LAbrini (Figure 3C).

**Figure 3.**
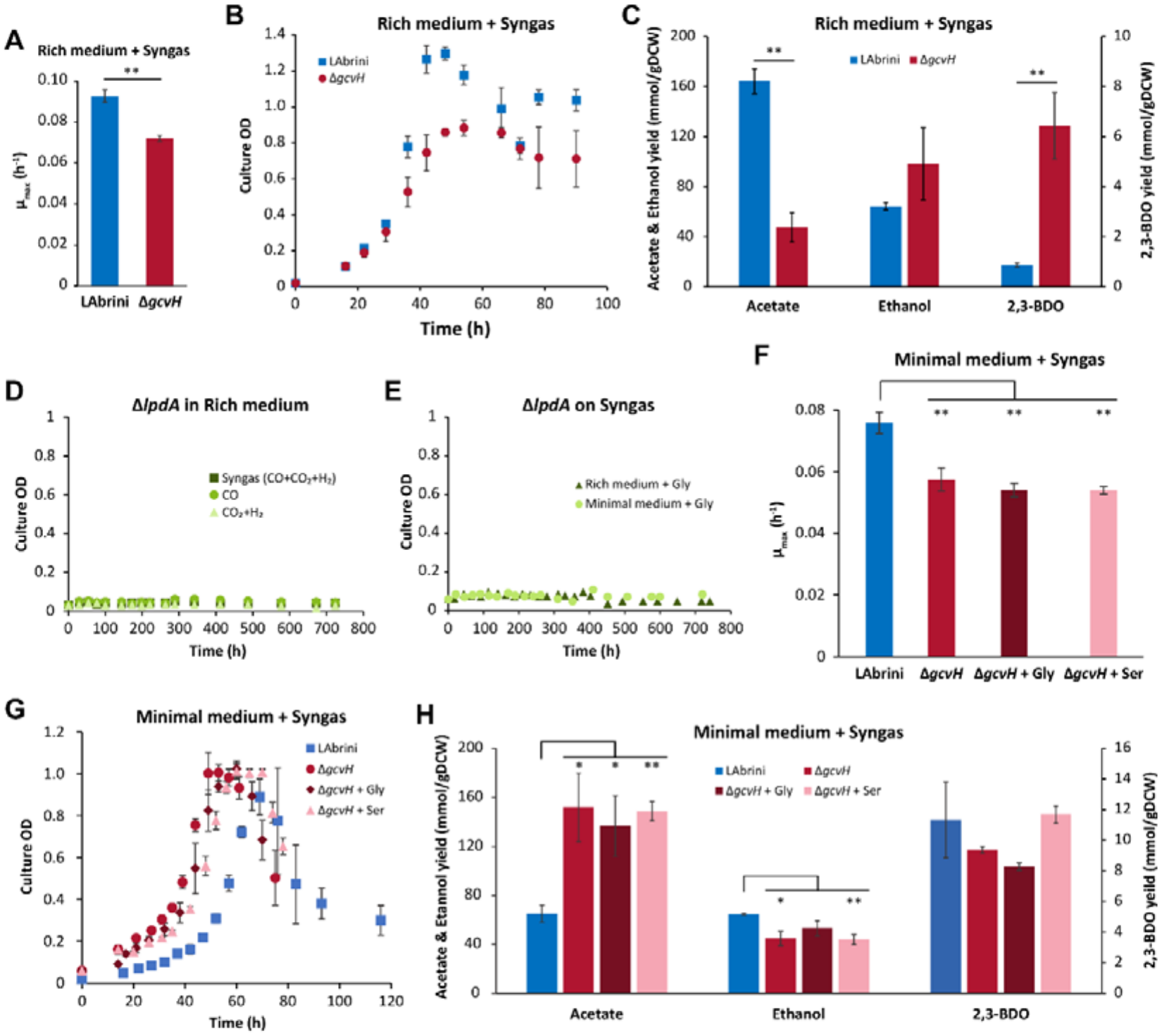
Functional and structural characterisation of the glycine cleavage system (GCS)-associated C_1_-fixing gene cluster genes *lpdA* and *gcvH* during autotrophic growth in bottle batch cultures. (A) Maximum specific growth rate (µ_max_) of LAbrini and Δ*gcvH* on syngas (CO+CO₂+H₂) in PETC-MES medium with YE (rich medium). (B) Growth profiles of LAbrini and Δ*gcvH* under the same conditions as in (A). (C) Growth by-product yields of LAbrini and Δ*gcvH* under the same conditions as in (A). (D) Autotrophic culture profiles of Δ*lpdA* on syngas (CO+CO₂+H₂), CO or CO₂+H₂ in rich medium. (E) Autotrophic culture profiles of Δ*lpdA* on syngas with 55 mg/L glycine (Gly) supplementation in rich medium or PETC-MES medium without YE (minimal medium). (F) µ_max_ of LAbrini and Δ*gcvH* on syngas in minimal medium, with or without Gly or 54 mg/L serine (Ser) supplementation. (G). Growth profiles of LAbrini and Δ*gcvH* under the same conditions as in (F). (H) Growth by-product yields of LAbrini and Δ*gcvH* under the same conditions as in (F). Data represent mean ± standard deviation between three biological replicates, except for Δ*lpdA* autotrophic growth and glycine supplementation cultures (single replicates). Asterisk denotes values statistically different according to two-tailed unpaired t-test (*, p < 0.05; **, p < 0.01). LAbrini, *C. autoethanogenum* LAbrini strain; YE, yeast extract; 2,3-BDO, 2,3-butanediol.

In contrast, Δ*lpdA* showed no growth on syngas, CO, or CO_2_+H_2_ (Figure 3D), providing the first experimental evidence of its essentiality for autotrophic growth, as essentiality for autotrophy could not be evaluated for genes essential for heterotrophy in the *C. autoethanogenum* TraDIS study (e.g. *lpdA*).^38^ The same study identified *gcvH* as essential for growth on CO in minimal medium. However, our targeted deletion results suggest that although both proposed GCS-associated genes are highly conserved within the C_1_-fixing gene cluster in acetogens,^16,17^ only *lpdA* is essential for *C. autoethanogenum* growth on syngas and may therefore constitute a core component supporting WLP function.

### 3.3 Glycine and serine supplementation effects on Δ*lpdA* and Δ*gcvH* autotrophic growth

As both LpdA and GcvH are proposed to encode GCS-associated proteins linked to WLP function during autotrophic CO_2_ fixation,^16^ we next investigated whether impaired glycine cleavage could account for the autotrophic defect observed in Δ*lpdA* and Δ*gcvH*. Both strains were thus cultivated in PETC-MES medium without YE (hereafter, minimal medium) in autotrophic bottle batch cultures, and Δ*lpdA* cultures were supplemented with glycine, whereas Δ*gcvH* cultures received glycine or serine, as glycine can be reversibly converted to serine via serine hydroxymethyltransferase to potentially bypass impaired GCS flux.^41^ Initial tests with 5.5 mg/L glycine and 5.4 mg/L serine, estimated to support synthesis of 0.23 gDCW/L biomass (OD = 1),^20^ did not support growth (data not shown); therefore, 10-fold higher concentrations (55 and 54 mg/L; respectively) were used for subsequent experiments. Data for LAbrini on syngas with minimal medium are from a previous study.^19^

Notably, glycine supplementation failed to restore autotrophic growth of Δ*lpdA,* even in rich medium (Figure 3E), indicating that this growth defect is not attributable to glycine limitations alone. This aligns with a broader role for *lpdA* in sustaining WLP-associated metabolism through its function in the GCS, where it oxidises reduced lipoyl intermediates and generates NAD(P)H to support redox-dependent metabolism.^42^ In acetogens, flux through the WLP relies on tightly coordinated NADH-, NADPH-, and Fd-dependent reactions, with methylene-THF serving as the key C_1_ intermediate in the methyl branch.^43^ Accordingly, loss of *lpdA* would be expected to perturb both reducing-equivalent availability and methylene-THF-linked C_1_ reduction reactions, thereby constraining methyl-branch flux during autotrophic growth. This interpretation is consistent with genome-scale metabolic modelling of *C*lostridium *drakei*, which predicts functional cooperation between glycine-associated pathways and the WLP to balance redox during autotrophy.^16^ It also aligns with recent work in the closely related *Clostridium ljungdahlii*, where LpdA was proposed to function primarily in GCS-associated methylene-THF and NADH metabolism.^44^ Notably, disruptions in redox balance and C_1_ metabolism can strongly reduce metabolic robustness of *C. autoethanogenum* during autotrophy.^36,45^ Thus, external supply of glycine or serine may not be expected to rescue such metabolic defects.

Similarly, supplementation with glycine or serine did not improve the growth of Δ*gcvH*, as µ_max_ values were lower than those of LAbrini, even in minimal medium (Figure 3F-G). Interestingly, unlike the phenotype observed in rich medium, Δ*gcvH* in minimal medium exhibited ∼2-fold higher acetate yields (152 ± 28; 136 ± 25 for +glycine; 148 ± 8 mmol/gDCW for +serine) than LAbrini (65 ± 7 mmol/gDCW) (Figure 3H). Although 2,3-BDO yields were unchanged, ethanol yields for Δ*gcvH* were ∼27% lower than LAbrini (65 ± 2 mmol/gDCW); indicating that deletion of *gcvH* alters carbon flux distribution without abolishing WLP functionality. This is in line with metabolic modelling predictions in which disruption of GCS-associated methylene-THF metabolism redirects carbon flux towards core WLP reactions while maintaining autotrophic growth.^46^

### 3.4 Computational analyses support specialised WLP-associated roles of LpdA and GcvH

To provide orthogonal evidence for the proposed functionalities of both genes, we complemented the experimental characterisation with SeqHub analysis of genomic context, protein domain annotation, structural prediction and protein interaction inference,^47^ together with Foldseek searches against experimentally characterised protein structures.Click or tap here to enter text.^48^ For LpdA, domain analysis identified the conserved FAD-binding (PF00890), pyridine nucleotide-disulphide oxidoreductase (PF13434), and NAD(P)-binding dimerisation (PF02852) domains characteristic of canonical dihydrolipoyl dehydrogenases (Figure S2). The predicted structure (average predicted Local Distance Difference Test [pLDDT] = 93) showed high structural similarity to the dihydrolipoyl dehydrogenase from *Elizabethkingia anophelis* (Template modelling [TM]-score = 0.953; 41% sequence identity; Table S1). These independent computational findings support the annotation of LpdA as a WLP-associated dihydrolipoyl dehydrogenase and are consistent with its essentiality for autotrophy in LAbrini (Figure 3).

The observation that *gcvH* deletion was non-lethal and that its phenotype was not driven by limitations in glycine or serine synthesis suggests that canonical GCS activity could be maintained through alternative means. Indeed, LAbrini encodes another GcvH-like protein (*gcvH2*; 02285), which is co-located in the genome with the canonical GCS genes *gcvPA*, *gcvPB*, and *gcvT* (Figure 4A), whereas the WLP-associated *gcvH* is conserved within the C_1_-fixing gene cluster as in other closely related acetogens,^16^ suggesting distinct physiological roles. Both proteins contain the conserved lipoylation lysine residue (Lys63; Figure 4B) and share only ∼60% nucleotide sequence identity (Figure S3), while *gcvH* retains the conserved GCV_H domain (PF01597; average pLDDT = 88) and shows high structural similarity to the GcvH protein from *Thermus thermophilus* (TM-score = 0.932). These findings suggest that *gcvH* performs a specialised lipoylation-associated role linked to WLP operation, while canonical glycine cleavage is maintained by *gcvH2*.

**Figure 4.**
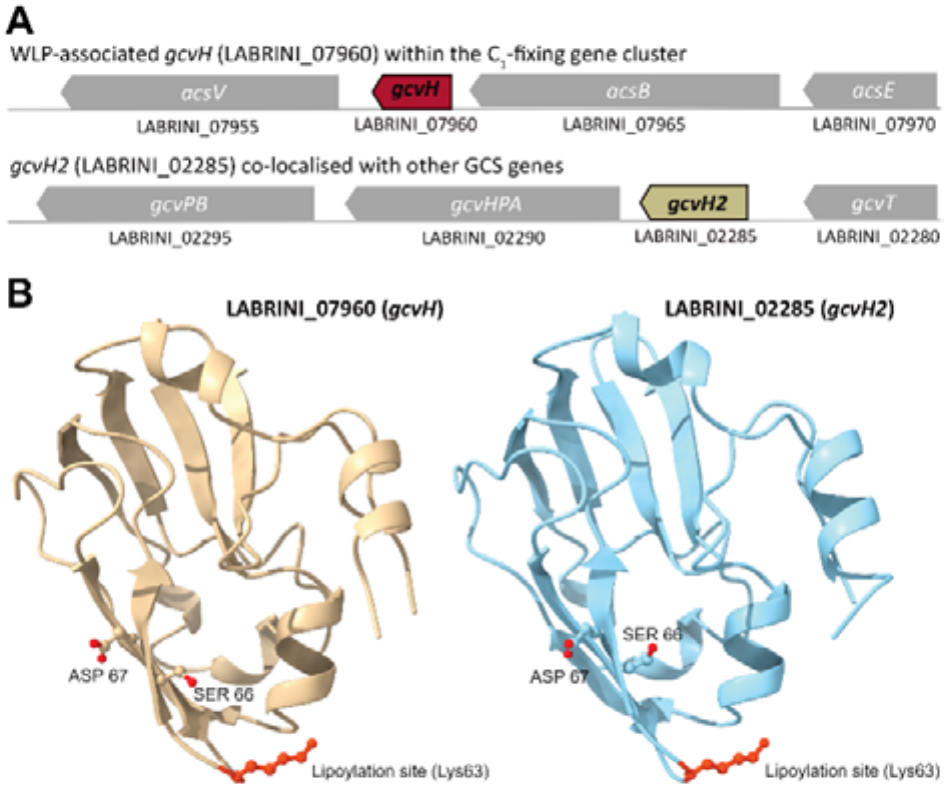
Comparison of the two GcvH-like proteins in *C. autoethanogenum* LAbrini. (A) Genomic organisation. (B) Structural comparison of SeqHub–predicted structures of GcvH and GcvH2, highlighting the conserved lipoylation site (Lys63) and neighbouring residues contributing to lipoyl-arm positioning.^40^ LAbrini, *C. autoethanogenum* LAbrini strain; WLP, Wood-Ljungdahl pathway; GCS, glycine cleavage system.

We note, however, that *gcvH2*, alongside the other canonical GCS genes, was classified as non-essential for autotrophy in the *C. autoethanogenum* TraDIS study.Click or tap here to enter text.^38^ Nevertheless, our findings are more readily reconciled with the genomic organisation described above and reports of functional diversification among GcvH-like paralogues in other bacteria,^49^ suggesting that *gcvH* fulfils a distinct role from *gcvH2*. Together, the *lpdA* and *gcvH* results indicate that neither gene functions primarily in glycine or serine biosynthesis, which is in line with previous modelling and ^13^C-labelling data,^46^ but instead, both contribute to autotrophic C_1_ and redox metabolism, supporting efficient WLP function.

### 3.5 *cooC* and *cooC2* are independently essential and have specialised roles in autotrophy

The second pair of genes – *cooC* and *cooC2* – have potentially linked but unclear functions within the C_1_-fixing gene cluster and are predicted to be MinD-type ATPases involved in CODH/ACS maturation.^29,31^ Whether the two proteins are functionally redundant or specialised remains unresolved. If it is the latter, deletion of either gene should independently abolish autotrophic growth, irrespective of whether it acts on CODH, ACS or both, since CODH/ACS activity is essential for autotrophy of *C. autoethanogenum*.^39^ Indeed, deletion of either *cooC* or *cooC2* abolished autotrophic growth in rich medium under all three tested gas conditions: syngas, CO, and CO_2_+H_2_ (Figure 5A), demonstrating that both genes are individually essential for autotrophy of LAbrini.

**Figure 5.**
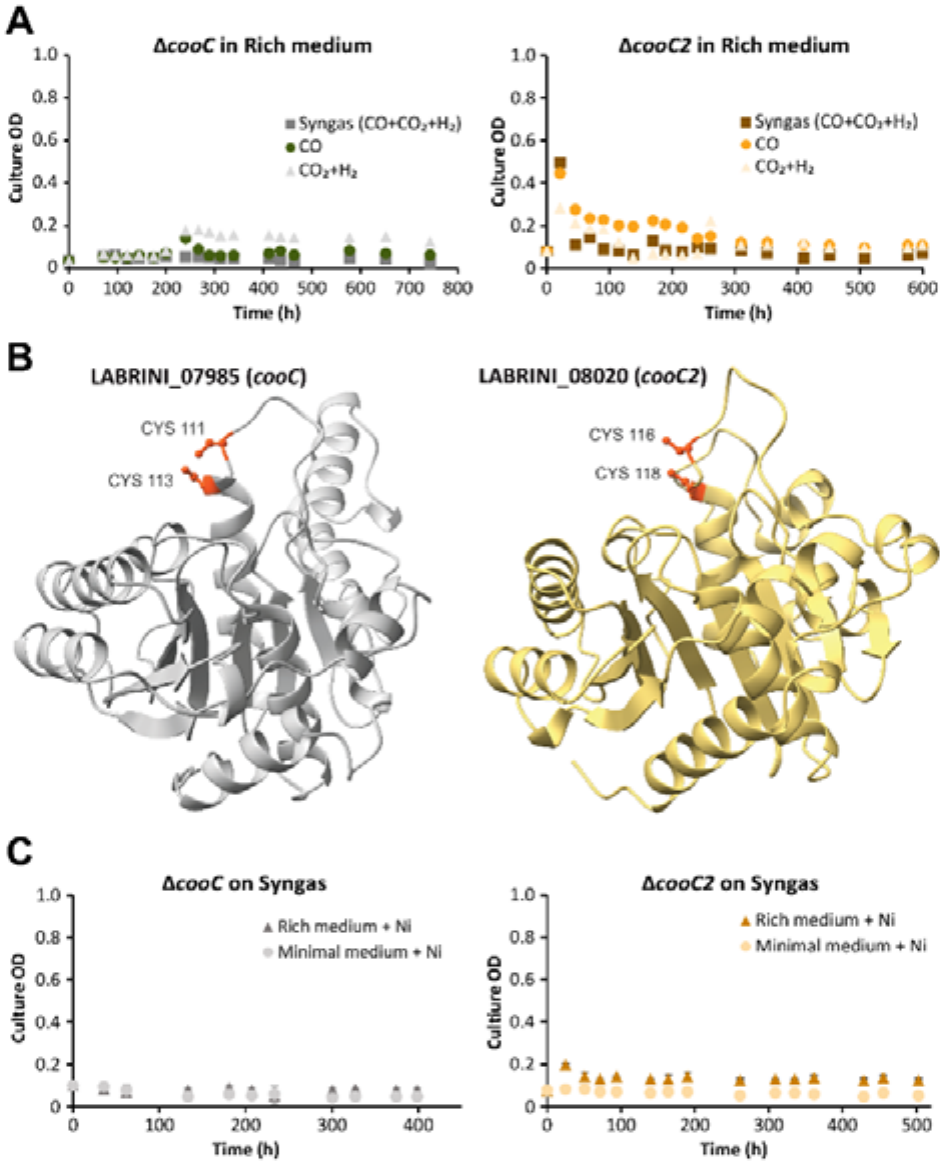
Autotrophic characterisation and nickel supplementation of Δ*cooC* and Δ*cooC2* in bottle batch cultures. (A) Growth profiles of Δ*cooC* and Δ*cooC2* on syngas (CO+CO₂+H₂), CO, or CO_2_+H₂ in PETC-MES medium with YE (rich medium). (B) SeqHub-predicted structures of CooC and CooC2 highlighting the Cys-X-Cys metal-binding motifs. (C) Culture profiles of Δ*cooC* and Δ*cooC2* on syngas with 0.8 mg/L Nickel (Ni^2+^) supplementation in PETC-MES medium with and without YE (minimal medium). Data represent mean ± standard deviation between two biological replicates, except for autotrophic growth experiments without Ni supplementation (single replicates). LAbrini, *C. autoethanogenum* LAbrini strain; YE, yeast extract.

Previous TraDIS analysis classified *cooC2* as non-essential for *C. autoethanogenum* JA1-1 autotrophy, while essentiality could not be determined for *cooC*.^38^ Our single-gene deletions therefore resolve the previously undetermined requirement for *cooC* and revise the non-essential assignment of *cooC2*, highlighting the value of a targeted approach for clarifying the roles of closely related genes and refining gene essentiality calls within this cluster. Our results mirror studies related to ATP-dependent Ni-maturation factors in other organisms: *C. hydrogenoformans* AcsF is required for ATP-dependent Ni incorporation and activation of apo-ACS, whereas *Rhodospirillum rubrum cooC* mutants accumulate Ni-deficient apo-CODH.^29,31,50–52^ These studies show that related ATP-dependent maturation factors can act on different components of the CODH/ACS complex.

Combined with our observations that both *cooC* and *cooC2* are essential for autotrophy but show differential heterotrophic growth phenotypes (Section 3.1), we hypothesise that the two proteins likely perform specialised roles in CODH/ACS maturation. We therefore examined whether this potential specialisation is reflected in structural differences between CooC and CooC2.^50,53^

### 3.6 Structural analyses revealed shared Ni-maturation features but distinct architectures in CooC and CooC2

Complementary protein domain and structural analyses revealed that both proteins indeed belong to the P-loop NTPase superfamily and contain the conserved AAA_31 and CbiA domains (Figure S2), characteristic of SIMIBI-class ATPases.Click or tap here to enter text.^54^ Structural analysis further identified the conserved Cys-X-Cys motif in both proteins (Figure 5B), consistent with ATP-dependent Ni coordination during CODH/ACS maturation.Click or tap here to enter text.^31,53,55^ However, while CooC retained a typical MinD/CooC-type ATPase architecture, CooC2 possessed an additional ArsA-type ATPase domain and exhibited broader similarity across multiple SIMIBI ATPase families. Notably, Foldseek analysis confirmed these architectural differences by identifying different structural homologues for the two proteins. The closest structural match for CooC was the FleN-like ATPase from *Pseudomonas aeruginosa* PAO1 (TM-score = 0.771), supporting conservation of the ATPase fold but not a Ni-insertion function. In contrast, CooC2 most closely resembles the Ni-maturation ATPase CooC1 from *C. hydrogenoformans* (TM-score = 0.801), providing structural evidence for a role inmetal-cofactor maturation. Taking also the *cooC* and *cooC2* deletion phenotypes into account, our data collectively indicate that CooC and CooC2 share the conserved structural features expected of ATP-dependent Ni maturation factors but likely perform different specialised roles in CODH/ACS maturation during autotrophic growth. Further work is needed to facilitate assignment of either paralogue to a specific metallocluster target (CODH C-cluster versus ACS A-cluster).

### 3.7 Nickel supplementation does not restore autotrophy of Δ*cooC* and Δ*cooC2*

Given the conserved ATPase architecture and Ni-coordinating motif identified in both CooC and CooC2, we next tested whether the autotrophic defects of Δ*cooC* or Δ*cooC2* reflected impaired Ni delivery to CODH/ACS. High Ni concentrations (500–650 µM) partially restore CODH activity and CO-dependent growth of *cooC* mutants in *R. rubrum* and *C. hydrogenoformans*.^50,51,56^ Thus, we tested if Ni^2+^ supplementation could alleviate the growth defect by cultivating Δ*cooC* and/or Δ*cooC2* on syngas in both rich and minimal media. Here, the defined Ni^2+^ concentration in our PETC-MES base medium was increased from 0.84 µM to 3.36 µM based on our previous Ni^2+^ supplementation study with LAbrini.^19^ We note that for the rich medium, 3.36 µM represents the minimum concentration because the Ni contribution from YE is undefined. Chelating agents used to alleviate Ni toxicity were avoided (e.g. casamino acids), because they could support growth and confound the interpretation of Ni^2+^ supplementation effects. Our concentration also exceeds the ∼1 µM residual Ni level, which is insufficient for realising CooC-dependent CODH/ACS activity in heterologous co-expression studies in *Clostridium acetobutylicum.*^52^ Notably, Ni^2+^ supplementation failed to restore autotrophy of either deletion strain (Figure 5C), indicating that increasing Ni availability, at least to ∼3.36 µM, is insufficient to compensate for the loss of either CooC or CooC2 for autotrophic growth of LAbrini.

These results suggest that the failure of Δ*cooC* and Δ*cooC2* to grow autotrophically is not attributable to insufficient physiologically available Ni, but rather to the disruption of ATP-dependent CODH/ACS maturation required for the assembly of its catalytically active metalloclusters. Furthermore, these data support growth test outcomes without additional Ni (Figure 5A), implying that CooC and CooC2 perform specialised, non-redundant functions during assembly of the bifunctional CODH/ACS complex.

### 3.8 AcsV is essential for autotrophic growth and shows features of an ATP-dependent corrinoid activase

CODH/ACS function depends not only on proper metal cofactor insertion but also on the corrinoid iron–sulfur protein (CFeSP), which transfers the methyl group from the methyl branch of the WLP to the CODH/ACS complex for acetyl-CoA synthesis.^9^ The last gene with an unclear function within the C_1_-fixing gene cluster – *acsV* – could be linked to corrinoid activation, as it is annotated to encode the acetyl-CoA accessory protein V, a corrinoid regeneration/activation protein associated with the bifunctional CODH/ACS complex.^14^ If AcsV solely fulfils this proposed role in *C. autoethanogenum*, deletion of *acsV* would interrupt corrinoid-dependent methyl transfer and thereby abolish autotrophic growth. Indeed, deletion of *acsV* abolished autotrophic growth in rich medium on syngas, CO, or CO_2_+H_2_ (Figure 6A). Similarly to the contradictions in gene essentialities between our work and the TraDIS study mentioned above, the latter classified *acsV* as non-essential for autotrophy.^38^ Overall, these discrepancies may reflect limitations of pooled transposon insertion fitness assays (e.g. TraDIS), including insertions falling within non-essential domains or individual fitness being masked in pooled populations.^57,58^

**Figure 6.**
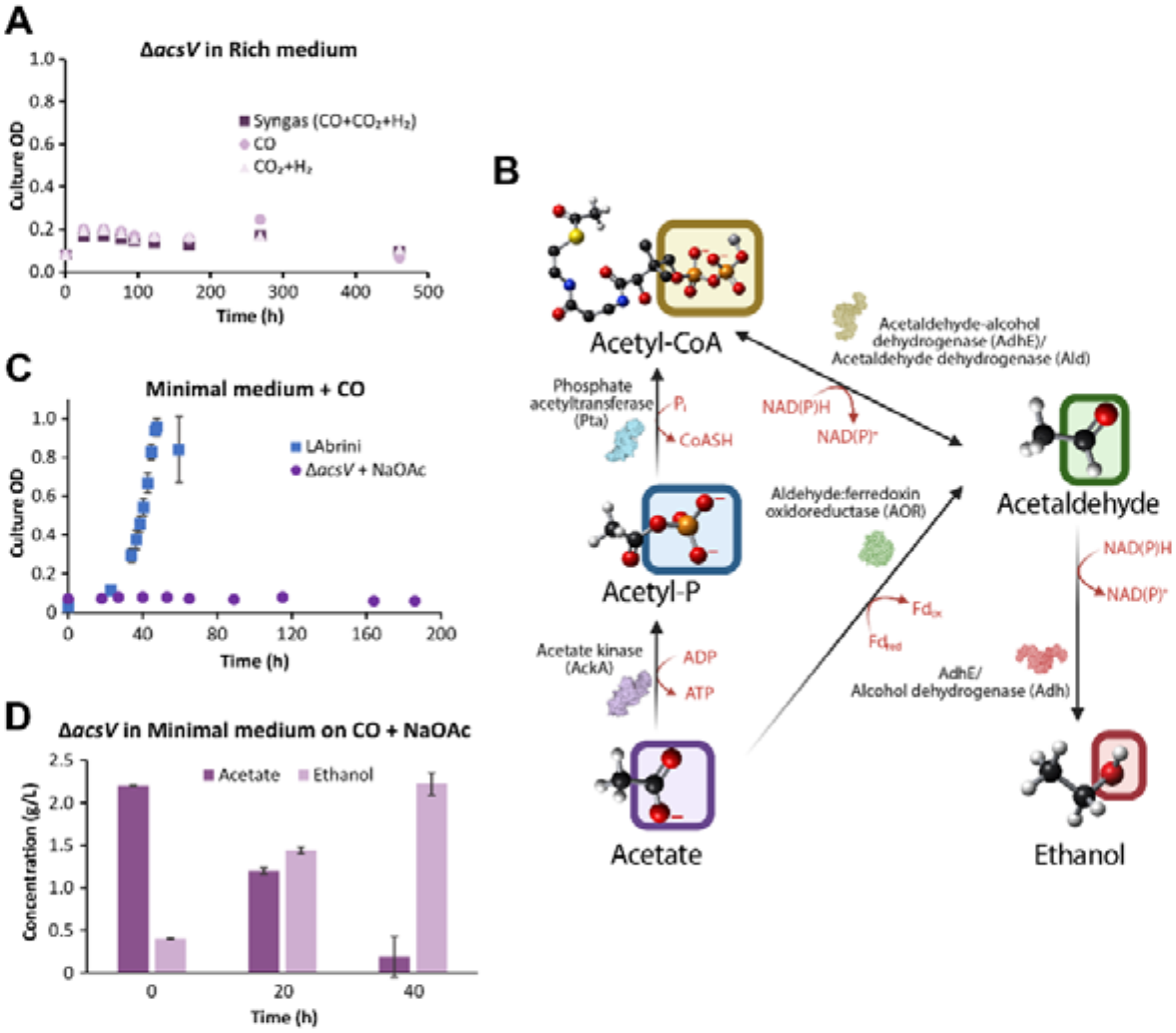
Autotrophic characterisation and acetate supplementation of Δ*acsV* in bottle batch cultures. (A) Autotrophic culture profiles of Δ*acsV* on syngas (CO+CO₂+H₂), CO, or CO₂+H₂ in PETC-MES medium with YE (rich medium) in bottle batch cultures. (B) Potential metabolic routes for acetate conversion to acetyl-CoA and ethanol in *C. autoethanogenum* (created using BioRender.com). (C) Culture profiles of LAbrini and Δ*acsV* on CO with sodium acetate (NaOAc) supplementation (2g/L) in PETC-MES without YE (minimal medium). (D) Time-course profiles of extracellular acetate and ethanol concentrations in Δ*acsV* cultures shown in (C). Data represent mean ± standard deviation between three biological replicates (error bars for Δ*acsV* on panel C are hidden by markers). LAbrini, *C. autoethanogenum* LAbrini strain; YE, yeast extract.

Within the WLP, the corrinoid cofactor bound to the CFeSP (AcsC/AcsD in *C. autoethanogenum*) cycles between the catalytically active Cobalt [Co(I)] and methylated Co (III) states during methyl transfer, and spontaneous oxidation to an inactive Co (II) periodically interrupts this cycle.^59^ Its reactivation requires dedicated ATP-dependent reductive activases of the RACE/Ram-family proteins, providing a specific benchmark against which candidate corrinoid activases can be evaluated.^60^ Domain and structural analyses identified conserved Fd, RACo linker, RACo middle, and ATPase domains in AcsV that are characteristic of ATP-dependent corrinoid activases (Figure S2). Foldseek further identified the RamQ corrinoid activase from *Carboxydothermus hydrogenoformans* as the closest structural homologue (TM-score = 0.897; Table S1) that belongs to the CFeSP-specific Ram/RACE subclade, rather than the broader group of activases acting on unrelated corrinoid-dependent methyltransferases, indicating functional as well as structural similarity. Protein interaction inference further identified AcsD, the corrinoid-binding CFeSP subunit, as the highest-confidence interaction partner of AcsV (confidence = 0.50) (Table S1). These complementary data support the annotation of AcsV as the ATP-dependent reductive activase responsible for maintaining the corrinoid cofactor in its catalytically active Co(I) state, making AcsV essential for autotrophic growth of *C. autoethanogenum*.

### 3.9 Acetate supplementation reveals that Δ*acsV* retains metabolic activity despite loss of autotrophy

If failure of corrinoid activation simply limited acetyl-CoA synthesis and would be the sole reason for lack of autotrophic growth, autotrophy in Δ*acsV* could be potentially at least partially restored by providing cells with non-glycolytic organic substrates from which acetyl-CoA could be readily produced by bypassing de novo acetyl-CoA formation. Assuming that Δ*acsV* cells can still oxidise CO, thereby generating reducing equivalents and ATP, supplying external acetate could enable acetyl-CoA production through acetyl-P or acetaldehyde (Figure 6B). We therefore supplemented Δ*acsV* cultures with sodium acetate (2 g/L) in minimal medium on CO, but still no growth was observed (Figure 6C). Notably, cells did, however, retain metabolic activity, as acetate was rapidly consumed and ethanol was produced (Figure 6D). These observations indicate that the inability of Δ*acsV* to grow on CO is not simply caused by limited acetyl-CoA synthesis, but instead reflects failure in core WLP functionality, likely through impaired corrinoid-dependent methyl transfer and, consequently, CODH/ACS catalysis.

### 3.10 Transcriptional repression of *acsV* alters autotrophic carbon partitioning but not growth

Lastly, we sought to further elucidate *acsV* function by testing the effect of its transcriptional repression on *C. autoethanogenum* autotrophic growth. To enable tunable and reversible gene repression,^61,62^ we constructed an inducible dCas9-based CRISPRi plasmid targeting *acsV* (pGFTi-*acsV*) and transformed LAbrini with it to generate the *acsV*_KD strain. A non-targeting plasmid (pGFTi-nTsgRNA) was similarly used to generate the control strain nTsgRNA_KD. Strains were grown on syngas in rich medium supplemented with 64 ng/mL anhydrotetracycline (aTc) to induce dCas9 expression; the control strain was also grown without the inducer to assess the effect of dCas9 expression on growth.

RT-qPCR data from exponential-growth samples showed 4-fold higher dCas9 expression in the induced nTsgRNA_KD cultures than in uninduced cultures (Figure 7A), confirming the functionality of the inducible promoter while also showing baseline promoter leakiness. Although dCas9 expression in *acsV*_KD was comparable to the uninduced nTsgRNA_KD control, it was still sufficient to strongly repress *acsV* expression by 6-fold (Figure 7B). Surprisingly, such significant repression of an essential gene had no effect on µ_max_ (0.052 ± 0.002 for *acsV*_KD vs 0.055 ± 0.002 for induced nTsgRNA_KD; p < 0.05) (Figure 7C). While *acsV* repression did not affect growth, the acetate yield was significantly reduced for *acsV*_KD without the corresponding increase in reduced by-product yields (Figure 7D). Thus, *acsV* repression altered autotrophic carbon partitioning without impairing growth.

**Figure 7.**
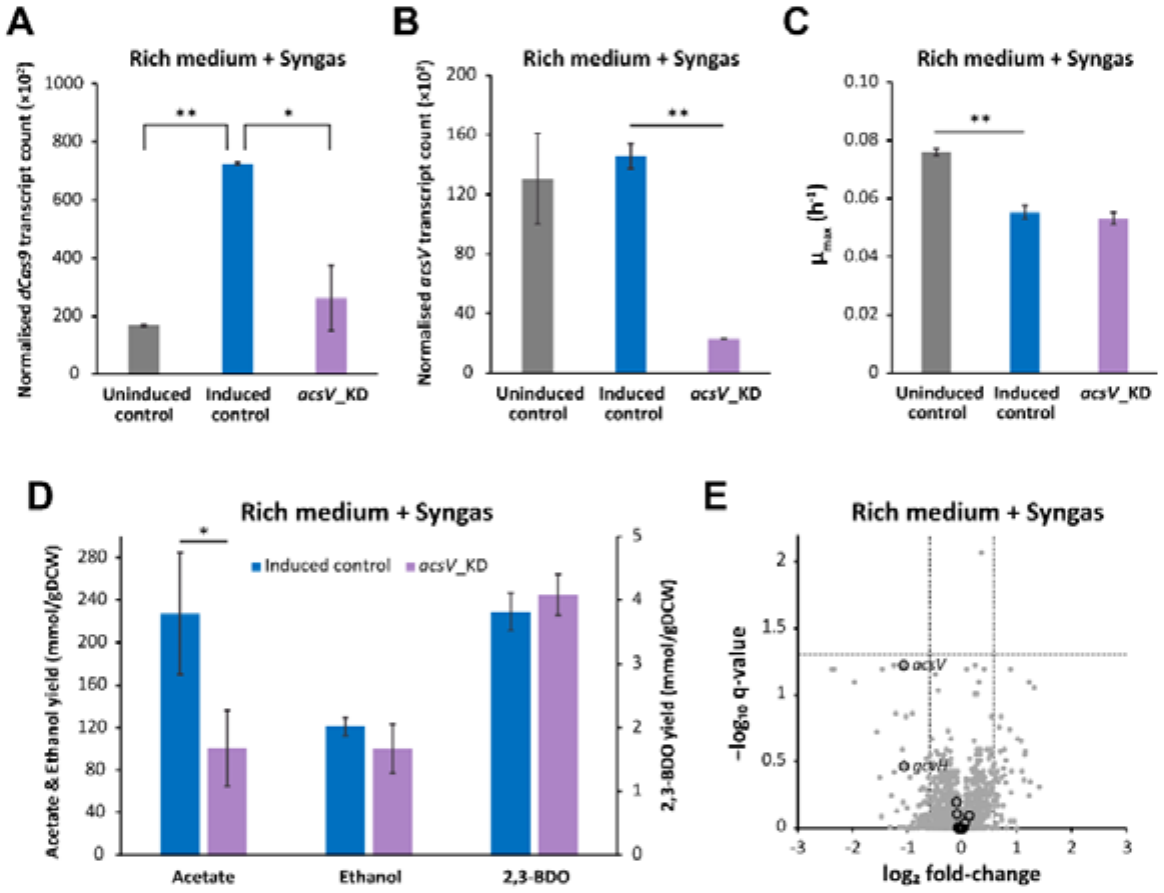
Autotrophic characterisation of *acsV* transcriptional repression in bottle batch cultures. (A) Normalised dCas9 transcript count (relative to 16S rRNA) from RT-qPCR analysis in the uninduced and induced non-targeting control strain (nTsgRNA_KD) and induced *acsV* knockdown strain (*acsV*_KD) grown on syngas (CO+CO₂+H₂) in PETC-MES medium with YE (rich medium). (B) Normalised *acsV* transcript count. (C) Maximum specific growth rates (μ_max_). (D) Growth by-product yields. (E) Volcano plot showing protein expression differences between *acsV*_KD and nTsgRNA_KD strains. Black open circles indicate proteins encoded within the C_1_-fixing gene cluster (LABRINI_07955–08021). Differential expression thresholds (fold-change > 1.5 and q < 0.05) are indicated by dashed lines. Data represent mean ± standard deviation between two biological replicates. Asterisk denotes values statistically different according to two-tailed unpaired *t*-test (**, p < 0.01; *, p < 0.05). KD, knockdown; gDCW, gram of dry cell weight; 2,3-BDO, 2,3-butanediol.

### 3.11 Unchanged proteome expression in *acsV*_KD explained by its translational buffering

To quantify the proteome response to acsV repression, we used proteomics on the *acsV*_KD and induced nTsgRNA_KD cultures described above. While we confidently detected expression of 2,005 proteins across the strains, surprisingly, none were differentially expressed between the two strains (fold-change (FC) > 1.5 and q-value < 0.05) (Table S2). Hence, AcsV abundance was not repressed – FC of −2 between *acsV*_KD and nTsgRNA_KD, but with q = 0.06 – at a 6-fold transcript repression (Figure 7E). Within the rest of the non-differentially expressed proteins from the C_1_-fixing gene cluster (07955‒08025), the adjacent GcvH had a decrease in abundance similar to AcsV (FC = −2; q = 0.34), while protein levels for others were indifferent between strains (FC < 1.1; q = 1.00). Importantly, the lack of AcsV repression (at protein level) explains why growth was not affected in *acsV*_KD. This can likely be attributed to translational buffering, in which changes in transcript levels do not necessarily translate into proportional changes in protein abundance due to differences in protein turnover, translational efficiency, and growth dilution.^63^ Still, acetate production was significantly affected in *acsV*_KD, suggesting sensitivity of CODH/ACS function to AcsV levels (Figure 7D). Notably, multi-omics analysis of *Escherichia coli* CRISPRi strains shows that even substantial reductions in enzyme levels do not necessarily impair cellular fitness; instead, they produce local metabolic rearrangements, demonstrating that metabolic networks can buffer enzyme perturbations while maintaining growth.^64^ Overall, *acsV*_KD data suggest that AcsV abundance affects acetate production before AcsV levels become limiting for *C. autoethanogenum* autotrophic growth.

## Conclusions

Targeted single-gene deletions demonstrated that *acsV, lpdA, cooC*, and *cooC2* are essential for autotrophic growth of *C. autoethanogenum*, whereas deletion of the WLP-associated *gcvH* reduced growth and altered carbon flux without abolishing autotrophy. Our physiological and supplementation data further show that *lpdA* and *gcvH* do not function primarily in glycine or serine biosynthesis, pointing instead to roles in WLP-linked C_1_ and redox metabolism, while the non-lethal Δ*gcvH* phenotype suggests functional divergence from the canonical glycine cleavage protein *gcvH2*. Structural analyses, together with the divergent deletion phenotypes observed here, support non-redundant roles for CooC and CooC2. In addition, acetate supplementation to Δ*acsV* revealed that metabolic activity is retained despite failure to restore growth, suggesting a core WLP defect due to *acsV* deletion, consistent with computational inference that AcsV is an ATP-dependent corrinoid activase. Together, our findings refine the functional organisation of the C_1_-fixing gene cluster in acetogens and support their metabolic engineering as industrial chassis for gas fermentation.

## Materials & Methods

## Strains, Growth Media, and Culture Conditions

*E. coli* strains, NEB Turbo and NEBExpress (New England BioLabs), were used as hosts for cloning, plasmid construction, and propagation. For transformant selection, *E. coli* strains were routinely cultivated in Lysogeny Broth (LB) or on LB agar plates supplemented with 100 μg/mL of ampicillin. The *C. autoethanogenum* strain LAbrini^14^ deposited in the German Collection of Microorganisms and Cell Cultures (DSM 115981) was used as the base strain for genetic engineering. The genetically engineered *C. autoethanogenum* strains generated in this work are listed in Table S3.

For *C. autoethanogenum* growth on solid medium, YTF agar containing 10 g/L YE, 16 g/L tryptone, 5 g/L fructose, and 15 g/L agar was used,^19,65^ supplemented with 4 μg/mL clarithromycin when needed. For growth in liquid cultures, conditions were the same as previously described.^19^ Briefly, batch experiments were carried out in 250 mL Schott bottles containing 50 mL of liquid medium in biological triplicates (unless stated otherwise), incubated horizontally at 37°C with orbital shaking at 120 RPM under strictly anaerobic conditions. Cells were grown in chemically defined PETC-MES medium,^20^ with or without 1.5 g/L YE (i.e. rich or minimal medium, respectively) and supplemented with 0.4 g/L of cysteine-HCl·H_2_O as the reducing agent. The carbon source was either fructose (5 g/L) or one of the gas mixes with headspace pressurised to 140 kPa: syngas (50% CO, 20% CO_2_, 20% H_2_, 10% Ar; AS Linde Gas), CO (60% CO, 40% Ar; AS Linde Gas), or CO_2_ + H_2_ (67% CO₂, 23% H₂, 10% Ar; AS Linde Gas).

Growth was monitored by measuring culture OD with frequent sampling during the exponential growth phase to determine μ_max_ and product yields. μ_max_ was calculated from 3-to-6 data points from the exponential phase by linear regression, yielding correlation coefficients (R^2^) > 0.94 between culture time and ln(OD). Product yields for acetate, ethanol, and 2,3-BDO (mmol/gDCW) were calculated from the exponential phase by linear regression with R^2^ > 0.90, except for acetate for Δ*acsV* and Δ*cooC* on fructose, LAbrini and Δ*gcvH* on syngas in rich medium, and Δ*gcvH* on minimal medium (R² = 0.78–0.88); ethanol and 2,3-BDO for Δ*gcvH* (R² = 0.76 and 0.87, respectively) on syngas in minimal medium, and 2,3-BDO for LAbrini on fructose (R² = 0.79). Data for LAbrini on syngas in minimal medium are from a previous study.^19^

Biomass concentration was determined by measuring OD and using the correlation coefficient between culture OD and DCW, established at 0.23, as described previously.^66^ Extracellular metabolome analysis from filtered broth samples was performed as previously described.^20^ We note that cells produced 2R,3R-butanediol.

Molecular Biology Techniques. Routine *C. autoethanogenum* genomic DNA extraction for PCR diagnostics was performed using the PureLink Genomic DNA extraction kit (Invitrogen™, Thermo Fisher Scientific). Plasmid DNA was extracted from *E. coli* using FavorGen FavorPrep™ Plasmid DNA Extraction Mini Kit (Favorgen Biotech Corp). All purified genomic and plasmid DNA fragments were quantified using the NanoDrop™ 1000 Spectrophotometer (Thermo Fischer Scientific). Colony PCR for initial routine screening to confirm plasmid presence and gene editing was performed using Hot FirePol Blend master mix (Solis BioDyne). PCR for amplification of DNA fragments used in sequencing, cloning, and confirmatory screening for plasmid presence and gene editing was performed using the Phusion™ High-Fidelity DNA Polymerase (Thermo Fisher Scientific). Primers used in this study (Table S4) were designed using the NetPrimer software (PREMIER Biosoft International) and synthesised by Integrated DNA Technologies. PCR products and DNA fragments were purified from agarose gels using the FavorPrep Gel/PCR Purification Kit (Favorgen Biotech Corp).

## Generation of Gene Deletion Strains

Single-gene deletion strains were obtained as previously described.^19^ Briefly, sgRNAs for targeted single-gene deletion of all five genes of interest were designed and cloned via Inverse PCR (iPCR) into a modified version of the pNickClos2.0 plasmid (Addgene #73228)^19,67^ and used as the backbone plasmid. Then, 1 kb of homologous regions upstream and downstream of each target gene were PCR-amplified and fused by overlap-extension PCR. The fused homologous region was then introduced into the corresponding intermediate sgRNA-containing plasmid via restriction cloning to obtain the final gene deletion plasmids containing all necessary parts for CRISPR/nCas9-mediated gene deletion: pGFT132 targeting *acsV* (07955), pGFT163 targeting *gcvH* (07960), pGFT151 targeting *cooC* (07985), pGFT161 targeting *lpdA* (07990), and pGFT131 targeting *cooC2* (08020) (Table S5). Next, electrocompetent LAbrini cells were electroporated with plasmids extracted from the Dcm^-^*E. coli* NEBExpress strain. Lastly, transformant colonies were screened by colony PCR and those with the targeted gene deletion were isolated for liquid outgrowth, cured of the plasmid using an optimised protocol,^68^ and glycerol stocked as five single-gene deletion strains of LAbrini: Δ*acsV*, Δ*gcvH*, Δ*cooC*, Δ*lpdA*, and Δ*cooC2* (Table S3).

## Generation of Gene Repression Strains

We used a previously developed in-house dCas9-based CRISPRi plasmid pGFT083,^69^ built on top of pNickClos 2.0,^67^ as the backbone for constructing the inducible CRISPRi system. For this, the constitutive pthl promoter in pGFT083 was replaced via overlap extension PCR with the aTc-inducible IPL12 promoter together with a TetR repression cassette,^70^ yielding pGFTi. This was then used as a template for iPCR to introduce a sgRNA for targeting *acsV* (pGFTi-*acsV*) and a sgRNA not targeting any gene in LAbrini (pGFTi-nTsgRNA) as a control plasmid. Next, plasmids were prepared and electroporated into LAbrini as described above. Lastly, transformant colonies were screened by colony PCR and those carrying the desired plasmid were isolated for liquid outgrowth and glycerol stocked as single-gene repression strains of LAbrini: *acsV_KD* and nTsgRNA_KD.

## Computational Protein Sequence and Structure Analysis

Protein amino acid sequences extracted from NCBI were analysed using SeqHub from Tatta Bio, accessed on 17.07.2026.^47^ Predicted functional annotations were derived from the closest Swiss-Prot homologues identified by SeqHub and interpreted alongside conserved domain architecture, genomic neighbourhood conservation, and predicted protein-protein interaction networks. Conserved domains were identified using the integrated domain annotation module in SeqHub. Genomic neighbourhood analysis was used to assess conservation of operon organisation and co-localisation with known components of the WLP based on the LAbrini genome annotation NCBI Genbank CP110420.^14^ Protein-protein interactions were predicted by the FlashPPI framework^71^ and only high-confidence interactions (prediction confidence ≥0.50) were considered. Three-dimensional protein structures were predicted using the integrated ESMFold2 framework^72^ and exported as Protein Data Bank (PDB) files. Average pLDDT confidence scores provided by ESMFold2 were used to assess overall model reliability. Predicted structures were subsequently compared against experimentally characterised proteins using FoldSeek,^48^ with default search parameters to identify the closest structural homologues. Conserved catalytic residues, metal-binding motifs, and other structurally relevant features were inspected by comparison with the highest-scoring structural matches. Finally, protein functional assignments were interpreted only when supported by multiple independent lines of evidence, including sequence annotation, conserved domain architecture, genomic context, structural similarity, and predicted protein-protein interactions. UCSF ChimeraX (version 1.9)^73^ software was used for the visualisation, organisation, and manipulation of the final predicted structures from SeqHub. The nucleotide sequences of *gcvH* and *gcvH2* were compared using Geneious Prime (version 2026.1.2; Dotmatics).^74^

## RT-qPCR Analysis

Transcript levels were quantified using RT-qPCR. For this, 15 mL of two biological replicate cultures of *acsV*_KD and nTsgRNA_KD were harvested during exponential growth (OD = 0.3‒0.4) with sample processing and total RNA preparation as described previously.^75^ Total RNA concentration was measured using the NanoDrop^TM^ 1000 instrument and 500 ng/20 μL was used as input for cDNA synthesis using RevertAid First Strand cDNA Synthesis Kit (Thermo Scientific) with random hexamer primers provided by the manufacturer.

RT-qPCR was performed on the LightCycler® 480 II Instrument (Roche) using the HOT FIREPol® EvaGreen® qPCR Supermix (Solis BioDyne) and custom primers for target genes (Table S4). The cycle threshold (C_t_) values for *dCas9* and *acsV* were normalised within biological replicates by the C_t_ value from *C. autoethanogenum* 16s rRNA genes (LAbrini_03960, 09860, 10650, 11110, 11180, 11255, 13590, 13740, and 17965; qPCR primers bind to all nine genes) and these normalised C_t_ values were used to quantify *dCas9* and *acsV* expression differences using the E^‒ΔΔCt^ method.

## Proteome Analysis

Proteome analysis comparing *acsV*_KD and nTsgRNA_KD was performed from two biological replicate cultures harvested during exponential growth (OD = 0.3–0.4). Sampling and sample preparation up to the determination of protein concentration in the lysates were performed as described previously.^19^ Then, 10 µg of protein was precipitated with trichloroacetic acid-deoxycholate (TCA-DOC) and dissolved in a denaturing buffer containing 7 M urea, 2 M thiourea, 100 mM ammonium bicarbonate, and 20 mM methylamine. Next, samples were reduced with 10 mM dithiothreitol for 30 min at 30°C and alkylated with 30 mM chloroacetamide in the dark for 30 min. Proteolytic pre-digestion was carried out with Lys-C (Fujifilm Wako Pure Chemical) at an enzyme-to-substrate ratio of 1:100 for 2 h at 25°C. The solution was diluted fivefold with 100 mM ammonium bicarbonate, and a second digestion was performed overnight at 25°C with dimethylated trypsin (Sigma-Aldrich) at a 1:50 enzyme-to-protein ratio. The produced peptide samples were acidified with trifluoroacetic acid (TFA) to a final concentration of 1.0% and desalted using custom-packed C18 solid-phase extraction tips (3M). Purified peptides were reconstituted in 0.5% TFA prior to nanoLC-MS/MS analysis using data-independent acquisition (DIA) followed by DIA MS data analysis as described before.^19^ The protein sequence database of LAbrini from NCBI GenBank (CP110420) was used. We confidently quantitated 28,178 peptides and 2,005 proteins across all samples, and 26,126 peptides and 1,874 proteins on average within each sample after removing shared peptides from analysis. Protein expression FC with q-values (after FDR correction)^76^ between *acsV*_KD and nTsgRNA_KD were determined using the software Perseus^77^ with Student’s T-test (Table S2) while only considering proteins quantified with at least two peptides (1,900 across all samples). Proteomics data have been deposited to the ProteomeXchange Consortium (http://proteomecentral.proteomexchange.org) via the PRIDE partner repository^78^ with the data set made public after manuscript publication.

## Author Information

### Author Contribution

Conceptualisation: U.J.N., K.R. and K.V.;

methodology: U.J.N, K.R. and K.V.;

formal analysis: U.J.N., M.V. and K.M.S;

investigation: U.J.N., M.V. and K.M.S.;

resources: K.V.;

writing – original draft: U.J.N. and K.V.;

writing – review and editing: U.J.N. and K.V.;

supervision: K.R. and K.V.;

project administration: K.V.;

funding acquisition: K.V.

## Conflict of Interest

Authors declare no conflict of interest.

### Supporting Information

Additional experimental details including results of computational analyses and predictions using SeqHub and FoldSeek; proteome analysis for differentially expressed proteins; lists of strains, primers, and plasmids (XLSX); results of Sanger sequencing for KO strains showing single-gene deletions; results of predicted protein domain architectures; and nucleotide sequence comparison of GcvH-like proteins (DOCX).

## Acknowledgements

We thank fellow group members Clara Vida G. C. Carneiro and Nilesh Kolhe for their feedback on the manuscript. This work was funded by the European Union’s Horizon 2020 grant agreement N810755 and the European Union (ERC, GENESYS, 101125746). Views and opinions expressed are, however, those of the authors only and do not necessarily reflect those of the European Union or the European Research Council. Neither the European Union nor the granting authority can be held responsible for them.

## Data Availability Statement

Proteomics data have been deposited in the ProteomeXchange Consortium (http://proteomecentral.proteomexchange.org) via the PRIDE partner repository,^78^ with the data set made public after manuscript publication.

## Notes

### Competing Interest Statement

The authors have declared no competing interest.

