## Supporting Figures for "Targeted Deletions Uncover Previously Uncharacterised C_1_-fixing Cluster Genes as Essential for Autotrophy in *Clostridium autoethanogenum*"

Ugochi Jennifer Nwaokorie^1^, Mariia Vorontsova^1^, Kurshedaktar Majibullah Shaikh^1^, Kristina Reinmets^1^, and Kaspar Valgepea^1, *^

^1^Institute of Bioengineering, University of Tartu, Tartu, Estonia

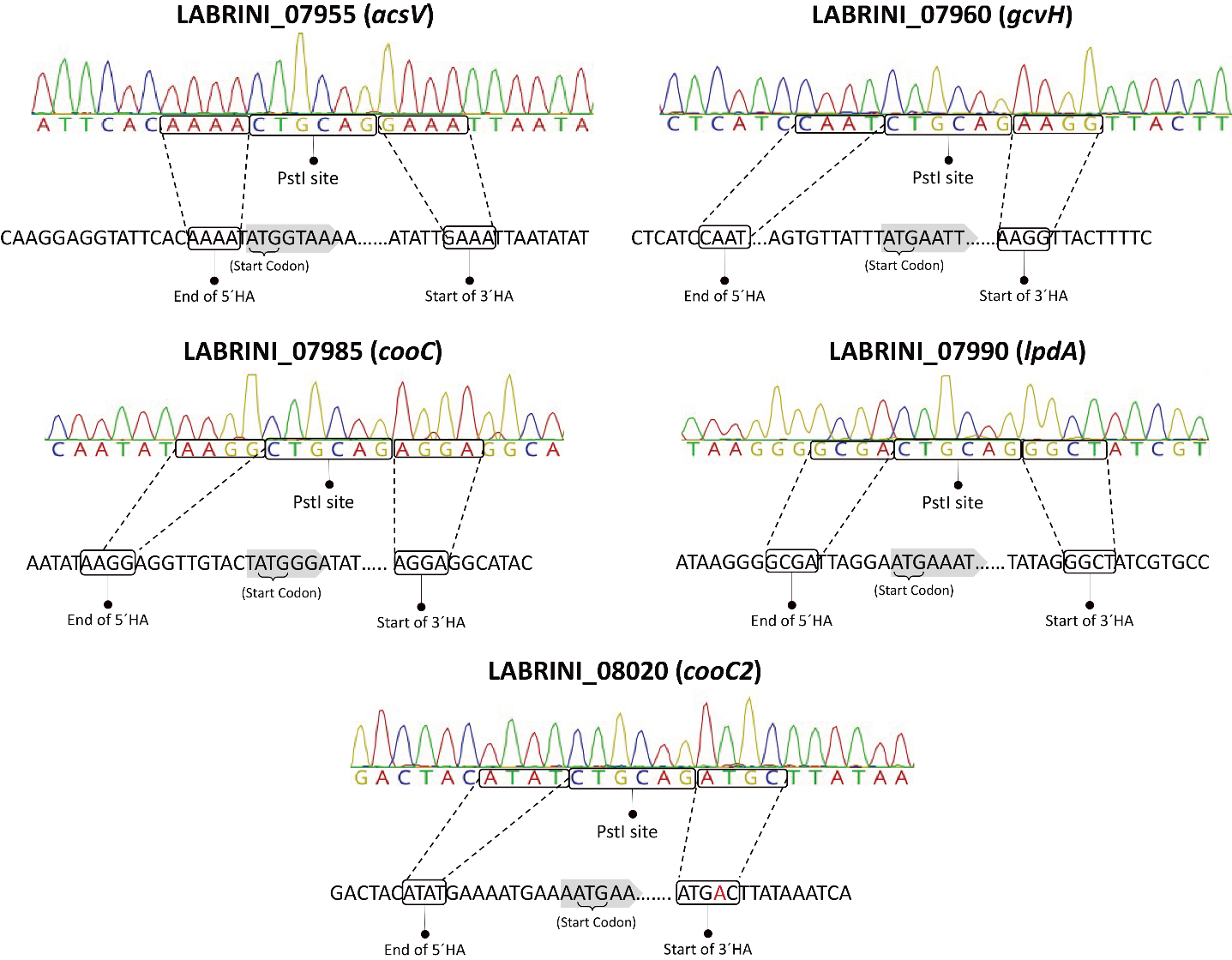

**Figure S1.** Sanger sequencing confirmation of gene deletions. HA, homology arm.

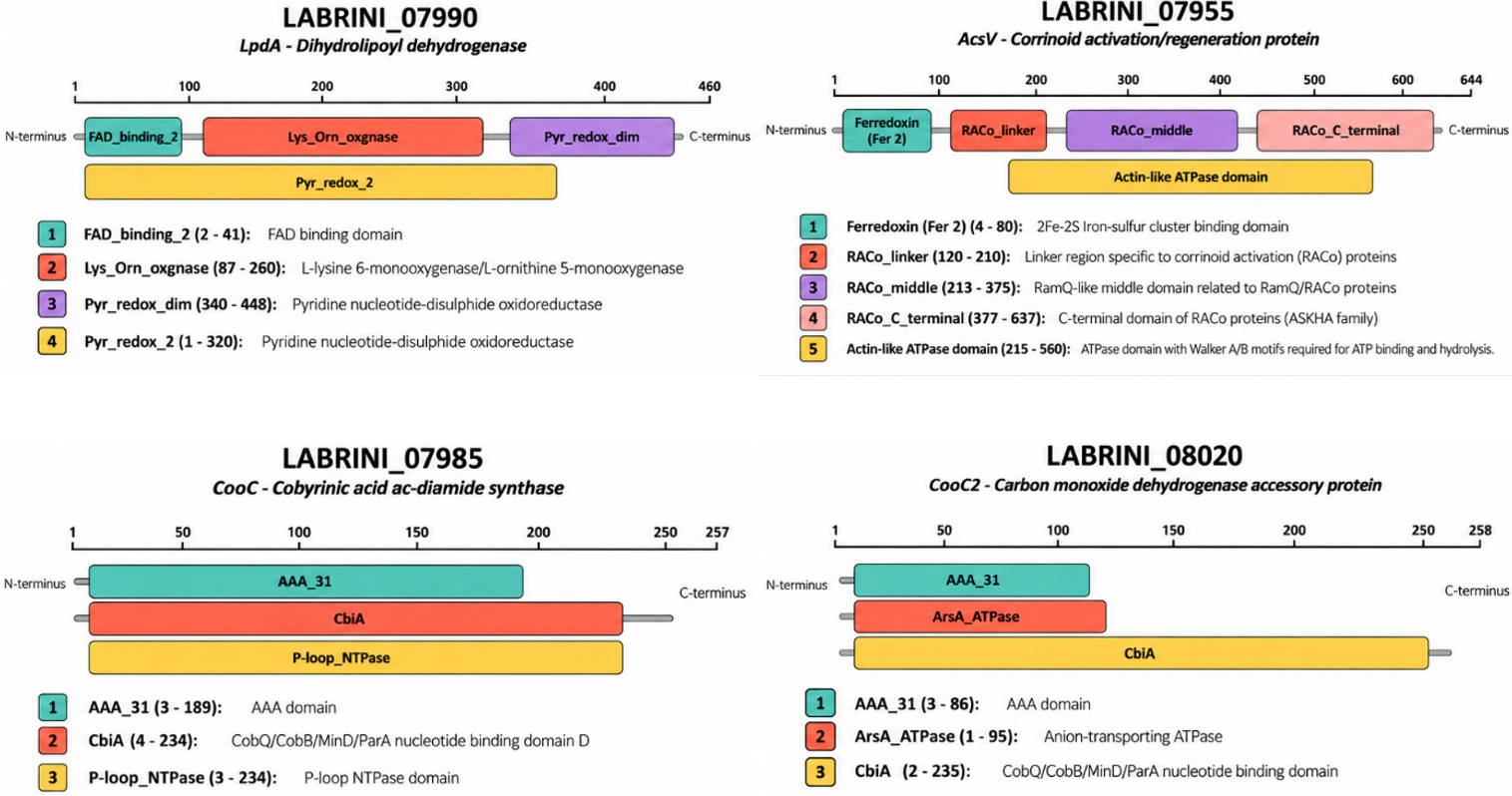

**Figure S2.** SeqHub-predicted protein domain architecture for AcsV, CooC, LpdA, and CooC2. See Materials and Methods in main text for details on SeqHub analysis. Results were visualised using OpenAI ChatGPT (GPT-5.6; OpenAI, 2026).

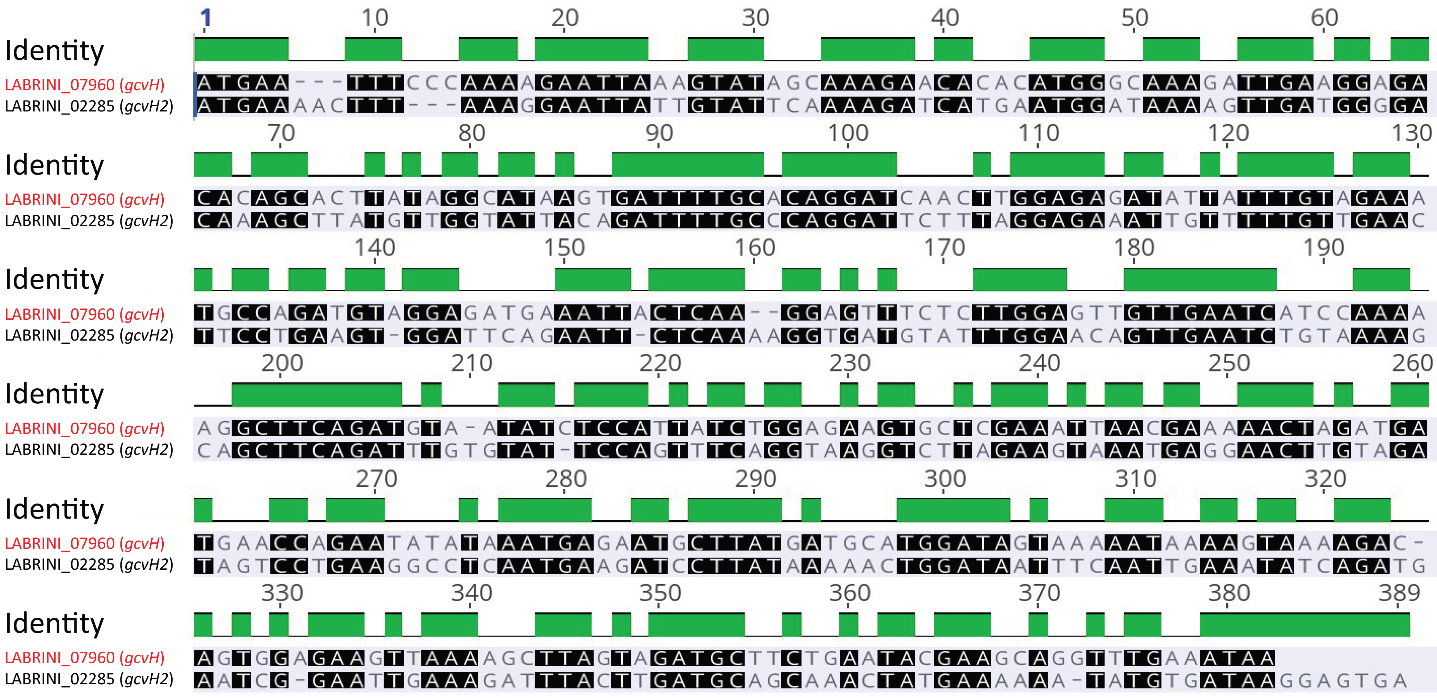

**Figure S3.** Sequence comparison of LABRINI_07960 (*gcvH*) and LABRINI_02285 (*gcvH2*) using the software Geneious. Green areas indicate identical nucleotides.
